# Inferring active cis-regulatory modules to predict functional regulatory elements

**DOI:** 10.1101/560862

**Authors:** Xi Chen, Andrew F. Neuwald, Leena Hilakivi-Clarke, Robert Clarke, Jianhua Xuan

**Author notes:** Corresponding author. Contact information: 900 N. Glebe Road, Arlington, VA 22203, USA.

## Abstract

Transcription factors (TFs) often function as cis-regulatory modules (CRMs) including both master factors and mediator coactivators to activate enhancers or promoters and regulate target gene transcription. Cell type-specific ChIP-seq profiling of multiple TFs makes it feasible to infer functional CRMs for a particular cell type. Yet, approaches based on co-localization of TF ChIP-seq peaks to infer CRMs are applied but many weak binding events, especially of those mediators, are missed by peak callers, resulting in an incomplete identification of CRMs. We developed a <u>ChIP</u>-seq data-based CR<u>M</u> inference approach with <u>G</u>ibbs-<u>S</u>ampling (ChIP-GSM). In a Bayesian framework, ChIP-GSM samples read counts of TFs iteratively for the joint effect of each potential TF combination. Using inferred CRMs as novel features, ChIP-GSM employs a logistic regression model to predict active regulatory elements. Performance validation on FANTOM5 enhancer or promoter regions revealed the superior performance of CRMs on regulatory region activity prediction than TFs. Finally, integrating CRMs inferred for K562 cells and gene expression data we found that CRMs are likely to activate regulatory regions or genes at different time points to mediate distinct cellular functions.

**Author Summary:** Accurately inferring cis-regulatory modules (CRMs) from a large set of TFs is a challenging task because the binding signals of TFs are often weak, noisy and sensitive to the cellular environment. Nevertheless, investigating TF associations may help understand the difference between enhancer and promoter activation mechanisms. In this paper, we develop a computational method (ChIP-GSM) to infer CRMs acting on regulatory elements at enhancer and promote regions. The novel method is built upon a Bayesian framework with Gibbs sampling that can be used to infer CRMs reliably hence to predict regulatory elements. The performance of ChIP-GSM is compared to that of existing methods, demonstrating its improved performance. Experimental results demonstrate that CRMs identified by ChIP-GSM are likely activating regulatory regions at different time points to mediate distinct cellular functions.

## 1. Introduction

Cis-regulatory modules are formed by binding ofhigh levels of master transcription factors and mediator coactivator [1, 2]. To mediate cell type-specific regulation, master transcription factors may recruit different coactivators in different context [3]. Given large-scale transcription factor (TF) ChIP-seq data in a particular cell type, joint analysis of multiple TFs binding sites co-localization has demonstrated power to recover cell type-specific CRMs [4, 5]. Yet, existing methods model are only applicable to a small number of TFs, i.e. master TFs, and the CRM inference is largely conditional on prior biological knowledge of TF associations. Given tens of TFs including coactivators and with little knowledge about their associations, existing methods cannot be directly used due to the model deficiency. Powerful CRM inference tools are needed to jointly model ChIP-seq profiles of a great number of TFs.

We developed a <u>ChIP</u>-seq data-based CR<u>M</u> inference approach with <u>G</u>ibbs-<u>S</u>ampling (ChIP-GSM). Given a regulatory region, we infer a CRM by sampling ChIP-seq read counts of multiple TFs iteratively for the joint effect of each potential TF combination on the current genomic location. Specifically, for each TF, we use a mixture model of Power-Law and Gamma distributions [6] to model read counts observed from binding or non-binding regions, respectively. Then, using Gibbs sampling, we infer the most reliable CRM (a combination of binding and non-binding TFs) at each regulatory region by sampling TF-associations and their read counts iteratively. After accumulating enough samples, ChIP-GSM estimates a probability for the occurrence of a candidate CRM at this region. We demonstrate ChIP-GSM’s ability to infer CRMs by applying it to ENCODE cell type-specific regulatory regions and TF ChIP-seq data. ChIP-GSM captures more regions regulated by high numbers of TFs. We also found that CRMs acting at enhancer and promoter regions are significantly different, suggesting that one should infer CRMs using one model for enhancers and another for promoters.

As regulatory regions are activated by CRMs having highly cell type-specific TF associations, using CRMs as features can more accurately identify cell type-specific active regulatory elements. Thus, ChIP-GSM employs a logistic regression classifier [7] to systematically predict the cell type-specific activity of each regulatory region. To validate the prediction accuracy, we combined ChIP-GSM results with prior knowledge of cell type-specific regulatory regions in the FANTOM5 database. The prediction performance of ChIP-GSM is significantly better than that of EMERGE [8] (a method featuring TF binding to predict regulatory activity). We studied CRMs inferred from promoter regions of K562 cells, which have the best prediction performance on active regions. Based on TF similarity between CRMs, we classified inferred CRMs from K562 promoter regions into several groups. We found that each group regulates very distinct sets of target genes. Further validation on time-course gene expression data of K562 cells shows that each group of CRMs are active at a specific time point and be involved in the regulation of a unique set of cellular processes. Therefore, ChIP-GSM can accurately infer biological meaningful CRMs, which provide strong prediction power on active cell type-specific regulatory elements.

## 2. Results

The flowchart of ChIP-GSM is shown in **Fig. 1**. Given ChIP-seq data of multiple TFs and candidate regulatory regions from the same context, we partition regions with different lengths into fixed-length (500 bps) bins and count normalized ChIP-seq read tags of every TF in each bin HOMER [9]. A PowerLaw model is fitted to the read counts of each TF at its binding regions (read count > 10 and fold change to the input signals > 2). A Gamma model is fitted to read counts at the same regions but from the matched input ChIP-seq profile. To infer CRMs, ChIP-GSM first identifies a list of candidate CRMs based on the frequency of TF co-colocalizations across all regulatory regions. Then, ChIP-GSM applies the probabilistic Power Law-Gamma mixture model to assign the weights of each TF to its binding or non-binding regions based on the observed ChIP-seq read count at each region. Under a Gibbs sampling framework, ChIP-GSM iteratively samples TF-associations, updates binding and non-binding regions of each TF, and recalculates weights of all TFs at each region. We run the sampling process until the sampler appears to converge on the equilibrium distribution. Then, ChIP-GSM accumulates samples of each assignment of a candidate CRM to a regulatory region and finally estimates a posterior probability (sampling frequency) representing the possibility of regulation. The inferred CRMs and their probabilities of regulation are further combined with experimentally measured cell type-specific regulatory regions in a supervised machine learning framework to predict the activity of every regulatory region. More details of the ChIP-GSM workflow can be found from ***Supplementary Fig. S1***.

**Figure 1.**
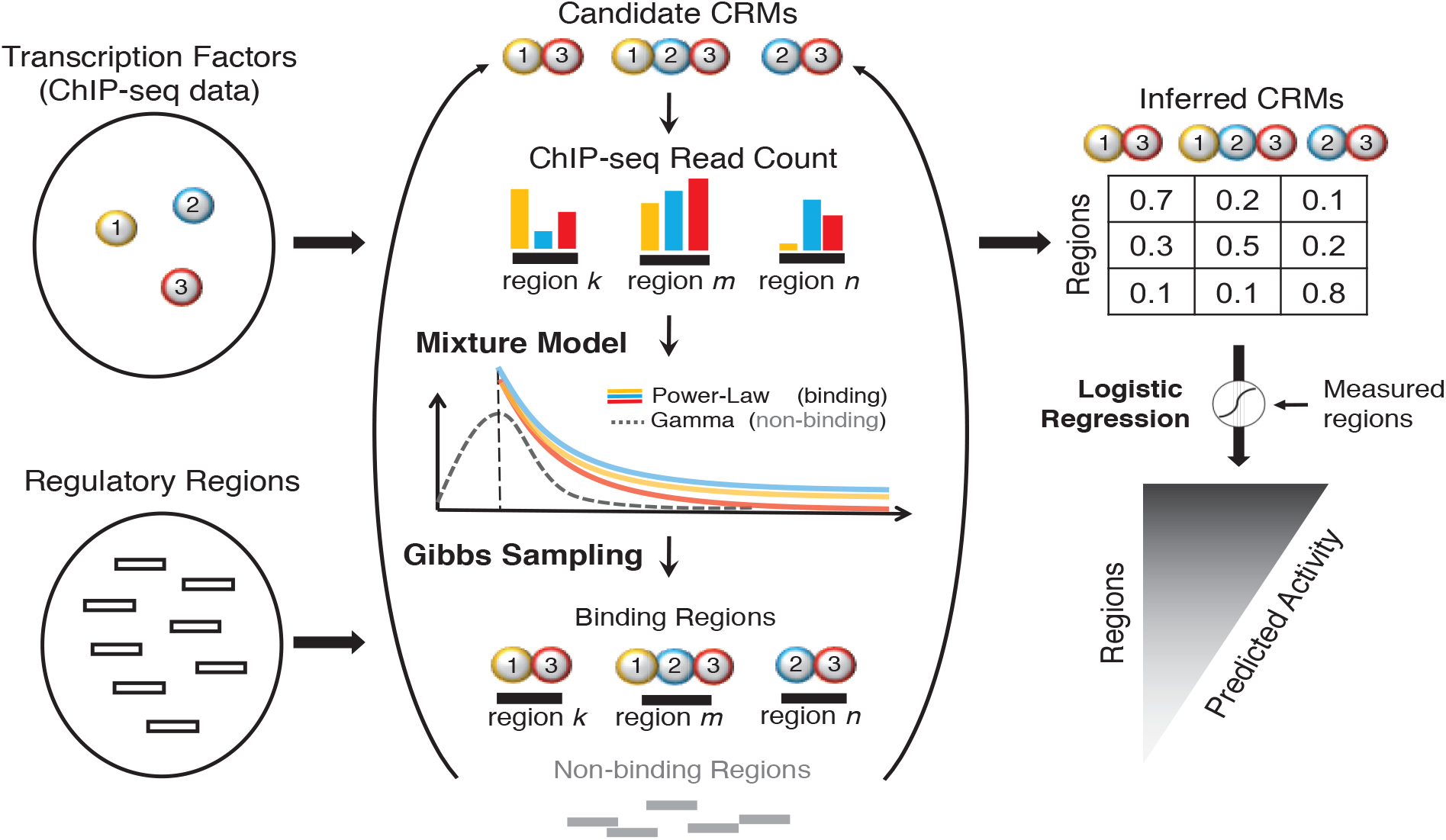
Flowchart of ChIP-GSM for CRM inference. Given ChIP-seq data of multiple TFs and candidate regulatory regions, ChIP-GSM features a mixture model of Power-Law (for binding events) and Gamma (for non-binding events) distributions to estimate read counts in binding and background regions. Using Gibbs Sampling, ChIP-GSM infers a CRM for each region in each round of sampling and finally estimates a posterior probability distribution of CRMs for each regulatory region. Integrating CRM probabilities with measured regulatory regions, ChIP-GSM further prioritizes all regulatory regions based on their activities predicted by a supervised machine learning approach.

### 2.1 CRMs function specifically at enhancer or promoter regions

An enhancer activator recruits other proteins binding at gene promoter regions to form a distal loop and initiates gene transcription [10]. TFs binding at enhancer regions can be different from proteins binding at gene proximal regions, then forming different CRMs. To examine these differences, we selected nine different cell types including breast cancer MCF-7, leukemia K562 and other major cell types. We downloaded their ChIP-seq data and cell type-specific enhancer- or promoter-like regions from the ENCODE database. Applying ChIP-GSM to the ChIP-seq data of each cell type, we respectively identified enhancer-CRMs and promoter-CRMs. Across all nine cell types, the similarity of CRMs between enhancer and promoter regions was 57% (**Supplementary Table S1**).

For MCF-7 cells, we identified 52 enhancer-CRMs and 66 promoter-CRMs, with a similarity of 54% (2×#common-CRMs/(#enhancer-CRMs + #promoter-CRMs)) (**Fig.2A**). For K562 cells, we identified 81 enhancer-CRMs and 67 promoter-CRMs, with a similarity of 55% (**Fig.2C**). For region-specific CRMs, we further checked the number of regions regulated by each CRM. As can be seen from **Fig.2B or Fig.2D**, in both MCF7 and K562 cell types, region-specific CRMs can be very strong and be regulating a high proportion of enhancer or promoter regions. We observed similar results from other cell types that have been tested. Thus, for the same set of TFs, their associations in CRMs between enhancers and promoters are different so it is necessary to respectively infer CRMs and further study their functions. ChIP-GSM identified enhancer or promoter CRMs of MCF-7, K562 and all other cell types can be found from **Supplementary Table S2**.

**Figure 2.**
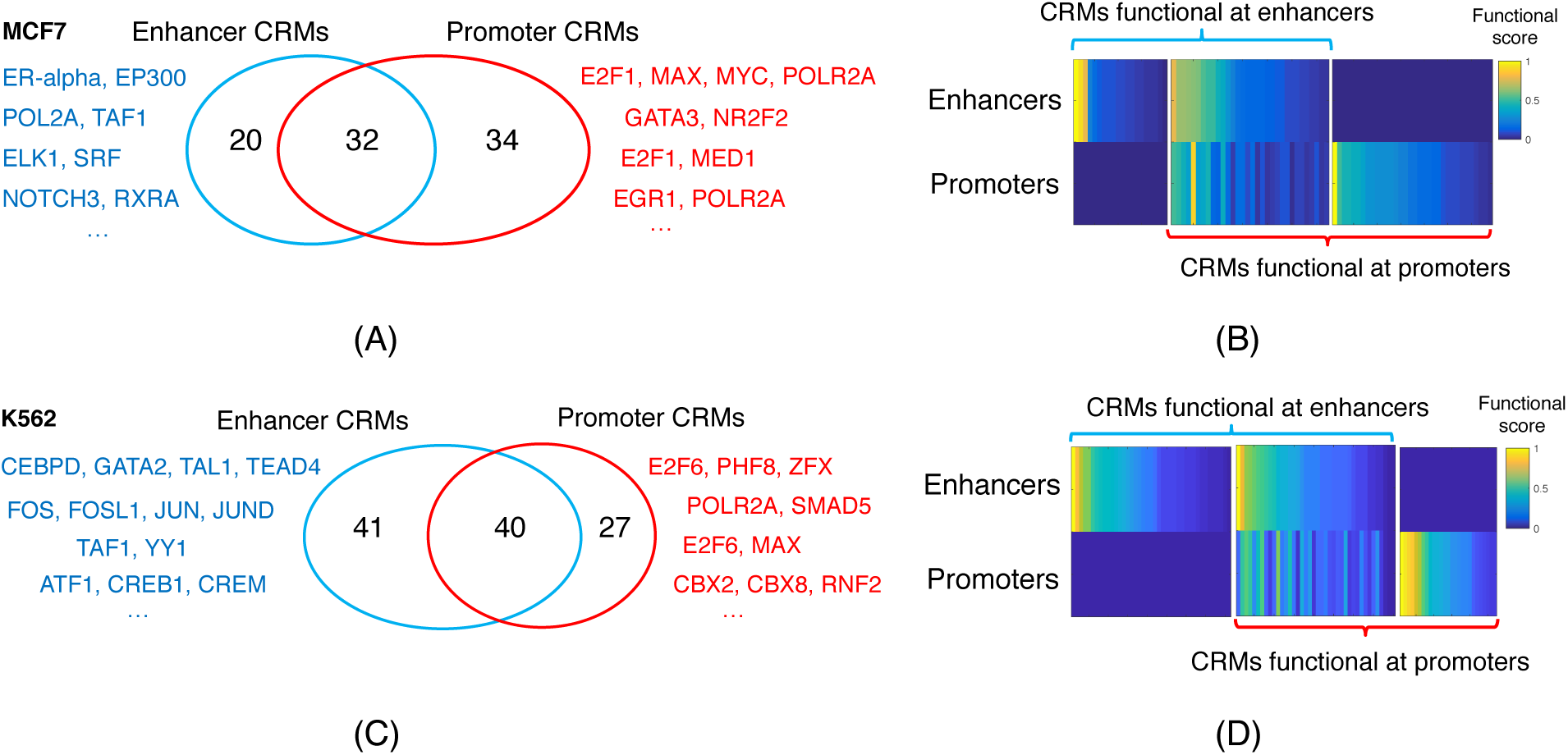
Inferred CRMs for enhancer and promoter regions respectively using ChIP-seq data of breast cancer MCF-7 cells. CRMs functioning at enhancer and promoter regions of (A) MCF-7 or (C) K562 cell types. Functional score of individual CRMs revealed that region-specific CRMs can be strong in (B) MCF-7 or (D) K562 cell types. Functional score is defined as the proportion of regulatory regions targeted by each CRM.

Specifically, for MCF-7 enhancer regions, the top two ChIP-GSM inferred CRMs are ERalpha-EP300 and POL2A-TAF1 (listed in **Fig.2A**). EP300 has been widely reported as a signature protein of enhancers. ER-alpha is a major enhancer activator in MCF-7 cells [11]. The association of TAF1-POLR2A was previously reported as an enhancer signature [12]. At gene promoter regions, the top CRM is E2F1-MAX-MYC-POLR2A. While MYC, MAX and POLR2A are frequently found in both enhancer and promoter studies of MCF-7 cells [13], E2F1 usually binds to gene promoters only [14]. This supports our finding of an association between a promoter specific factor E2F1 and a general module MAX-MYC-POLR2A only at gene promoters.

At K562 enhancers, ChIP-GSM reported a strong association of CEBPD-GATA2-TAL1-TEAD4. The overexpression of the TAL1 has already been reported being associated with super-enhancer formation in acute lymphoblastic leukemia. In the meanwhile, TEAD4 and CEBPD were both master transcription factors enriched at super-enhancers [2]. In pediatric acute myeloid leukemia, significant differences in survival were observed between patients with high versus normal GATA2 gene expression [15]. As a TF, GATA2 is prevalent at dynamic enhancers [16]. This association of CEBPD-GATA2-TAL1-TEAD4 was also found at K562 H3K4me1-enriched enhancer regions in another independent study [17]. At promoter regions, the top predicted module is E2F6-PHF8-ZFX. Similar to E2F1, E2F6 binding sites are demonstrated in a ChIP-chip study to be located proximal to TSSs, in both normal and tumor cells [18]. PHF8 binds to H3K4me3, an active histone mark enriched around TSSs [19]. ZFX was found in many tumor types by binding at CpG island promoters [20].

### 2.2 Predict enhancer or promoter activity using ChIP-GSM inferred CRMs

We further explore the predictive power of inferred cell type-specific CRMs on active regulatory regions. We used enhancer RNA (eRNA) or mRNA expression data from FANTOM5 (http://fantom.gsc.riken.jp/5/) as experimentally-based measures of cell type-specific enhancer or promoter activities. For each cell type, we labeled a region as ‘positive’ if it is active (TPM > 0.5 in at least two measures of this cell type) or ‘negative’ if it is non-active in the current cell type but active in at least one other cell type.

Combining ChIP-GSM results and labelled cell type-specific regulatory regions, we trained an elastic net logistic regression model to predict cell type-specific activity of each enhancer or promoter. For performance evaluation, each model was trained on 80% labeled regions and further tested on 20% hold-out regions. Numbers of labeled ‘positive’ or ‘negative’ regions as well as the training performance per cell type are listed in ***Supplementary Table S3***. The performance of ChIP-GSM featuring CRMs is significantly better than EMERGE [8], a competing model trained with TF binding events. Area under receiver operating characteristic curves (AUROC) were shown in **Fig. 3A** for enhancers (ChIP-GSM median AUROC 0.81 vs. EMERGE median AUROC 0.74; Wilcoxon rank-sum *p*-value < 0.05) and in **Fig. 3E** for promoters (ChIP-GSM median AUROC 0.83 vs. EMERGE median AUROC 0.73; Wilcoxon rank-sum *p*-value < 0.05).

**Figure 3.**
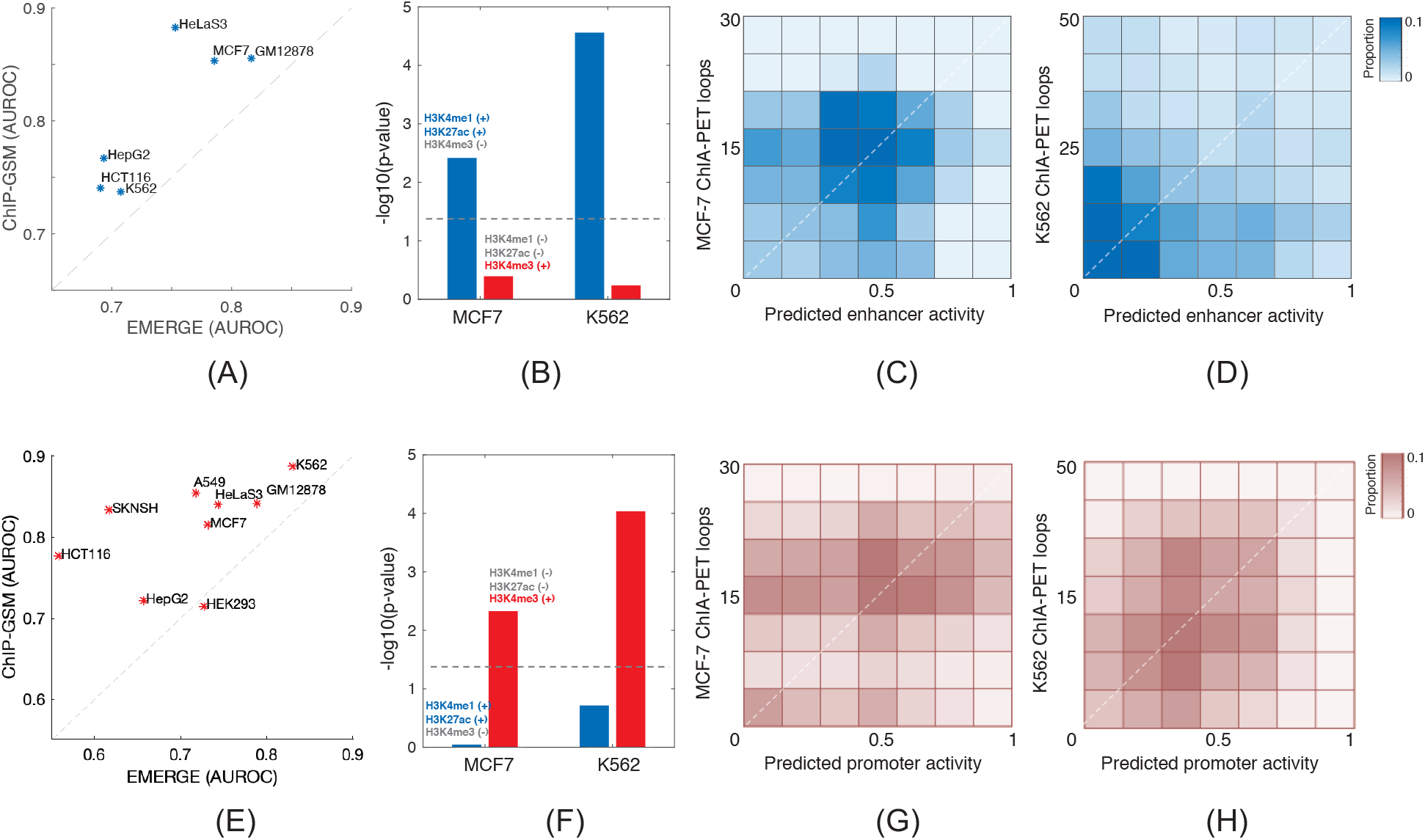
Prediction performance of active enhancers or promoters in nine different cell types using ChIP-GSM results. (A) ChIP-GSM predicted active enhancers more accurately than EMERGE. (B) The top predicted enhancers are significantly enriched with peaks bound by H3K4me1 and H3K27ac but not by H3K4me3. A significant correlation was observed between the predicted activities of enhancers and the number of enhancer-overlapping ChIA-PET loops in both MCF7 (C) and K562 (D) cell types. (E) Similarly, ChIP-GSM predicted active promoters more accurately than EMERGE. (F) The top predicted promoters are significantly enriched with peaks bound by H3K4me3 but not by H3K4me1 or H3K27ac. A significant correlation was observed between the ChIP-GSM predicted activities and the number of ChIA-PET loops around gene promoters in both MCF7 (G) and K562 (H) cell types.

To validate the prediction accuracy of enhancer or promoter activities, for MCF-7 and K562 cells, we first checked their enrichment of ENCODE cell type-specific ChIP-seq peaks of H3K4me1, H3K27ac and H3K4me3. H3K4me1 and H3K27ac are enhancer markers while H3K4me3 is a promoter marker [21]. As shown in **Fig. 3B**, in both MCF-7 and K562 cell types, the top 10% ChIP-GSM-predicted enhancer regions are significantly enriched with peaks with H3K4me1 (+), H3K27ac (+) and H3K4me3 (-) (hypergeometric p-values 2.76e-5 for K562 and 3.83e-3 for MCF-7). While in **Fig. 3F**, for promoters, the top 10% ChIP-GSM prediction are significantly enriched with peaks with H3K4me1 (-), H3K27ac (-) and H3K4me3 (+) (p-values 9.23e-5 for K562 and 4.7e-3 for MCF-7). The top predicted regions in one catalog are not significantly enriched with markers for the other catalog.

Moreover, enhancer or promoter activities are mediated by 3D chromatin interactions [22, 23]. Across all regions, we demonstrated that ChIP-GSM predicted regulatory activities are significantly correlated with the number of chromatin interactions in the matched cell type-specific ChIA-PET data in the ENCODE database (**Fig. 3;** p-value < 1e-10). The more active a regulatory region is predicted by ChIP-GSM, the higher number of chromatin interactions are observed around the same region. Therefore, through multiple studies we demonstrated that ChIP-GSM predictions are accurate and cell type-specific.

### 2.3 High-level CRM groups mediate distinct cellular functions

We finally explored CRM target genes and examine if there is function difference between genes that regulated by different CRMs. For this, we selected CRMs inferred from K562 cell promoter regions because: (1) these CRMs achieved the highest predictions of active promoters (**Fig. 3E**; AUC = 0.89); (2) compared to other methods capable of inferring CRMs from a large number of TFs ChIP-GSM identified more detailed and biologically important CRMs (***Supplementary Figs. S2-S5***). We selected in total 30 CRMs (with 39 TFs) where each CRM regulated at least 500 active promoter regions as predicted by the trained model (logistic regression score > 0.2).

We clustered these 30 CRMs based on the similarity of TFs between them using hieratical clustering and obtained eight groups (**Fig. 4A**; see ***Supplementary Fig. S2*** for details regarding each group). We found strong co-expression [24] (GEO access number: GSE1036) and coherent functions for TFs in each group (**Fig.4B** and **Supplementary Fig. S6**). In **Fig. 4B**, we show Groups 1, 3, 5, 6, 7 and 8, which contains at least one CRM with significantly co-expressed TFs (p-value<0.05, Fisher’s test). Although the overall co-expression in Group 2 is not significant, TFs in each group (CBX2, CBX8, RNF2, EZH2 and SUZ12) are all Polycomb-group proteins, an association that occurs in K562 cells [25]. Our results supported the hypothesis that cooperative TFs are usually active simultaneously [26, 27].

**Figure 4.**
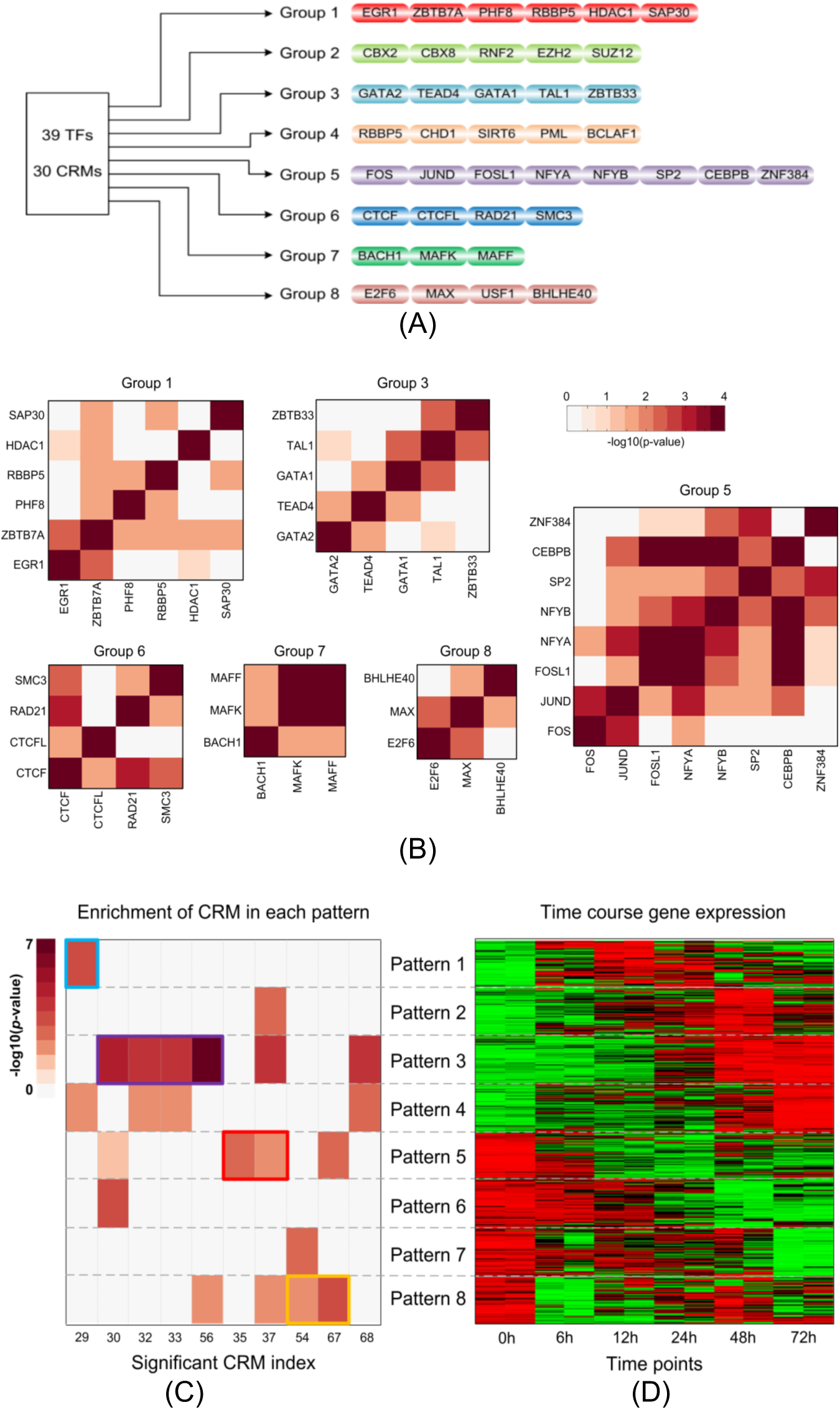
ChIP-GSM identified CRMs from the gene promoter region of K562 cells. **(A)** Eight groups of CRMs identified by ChIP-GSM functioning at gene promoter regions in leukemia K562 cell type; **(B)** mRNA co-expression of pairwise TFs in each group; **(C)** selected CRMs whose target genes are significantly enriched (hypergeometric p-value < 0.001) in active genes in a gene cluster in **(D)**. Each color label or box represents a unique CRM group.

For each CRM group, we did gene set functional enrichment using DAVID [28] for genes regulated by at least one CRM in that group. Results show that CRM groups tend to mediate different cellular functions (***Supplementary Fig. S7*** and ***Table S4***). For example, for genes regulated by CRMs in Group 1, acute myeloid leukemia was uniquely and significantly enriched; in Group 4, there were more genes involved in cancer development, including chronic myeloid leukemia; in Group 5, cell survival related functions including cell death, apoptosis, and p53 signaling pathway were uniquely enriched. These findings imply that genes co-activated by different sets of TFs are likely to involve in different cellular processes.

To further examine if genes regulated by different CRMs are active at different cell stages, based on gene expression, we clustered 1,569 differentially expressed genes in a K562 time-course gene expression dataset [24] to eight clusters using hierarchical clustering (**Fig. 4D**). Then, in each cluster, we assessed the enrichment of genes targeted by each CRM. Significant CRM-gene cluster pairs (hypergeometric *p*-value < 0.001) were shown in **Fig. 4C** (circled by ‘blue’, ‘purple’, ‘red’ and ‘yellow’ color, matching to the color code of CRM group in **Fig. 4A**) and enriched genes were listed in ***Supplementary Table S5***. CRMs from different groups are rarely functioning at the same time point. For instance, CRM 29 (labeled by the ‘blue’ box) is highly enriched in gene cluster 1. CRMs 30, 32, 33 and 56 (labeled by the ‘purple’ box) are all from Group 5 and their target genes are significantly enriched in gene cluster 3. CRMs 54 and 67 (labeled by the ‘yellow’ box) are very different and there is almost no overlap between their target gene lists. Nonetheless, CRM 54 and 67 are from module Group 4 and their target genes are enriched in gene cluster 8. These observations indicate that CRMs activate genes at different cell stages and may mediate different cell functions.

## 3. Discussion

Accurately inferring CRMs from a large set of TFs is challenging because, compared to histone marker, the number of TFs is much larger, and their binding signals are weaker, noisier, and more sensitive to the cellular environment. Nevertheless, investigating TF associations may help explain differences between enhancer and promoter activation mechanisms and target gene expression. In this paper, we developed a probabilistic model, ChIP-GSM, for inferring cell type-specific CRMs and for predicting active regulatory elements. ChIP-GSM better models ChIP-seq read counts for both strong and weak binding events, and thereby aids investigation of CRMs at regions containing weak binding signals that can be highly cell-type specific but easily missed by peak callers. ChIP-GSM currently infers CRMs at enhancer- or promoter-like regions. As a general computational framework, ChIP-GSM could be applied to inferring CRMs in any TF binding regions.

We demonstrated that CRMs are better than TFs at predicting active regulatory elements. Using K562 cells as an example, this study also reveals the functional diversity of CRMs: target genes of different CRMs are expressed at different time points and are involved in distinct cellular processes. We anticipate that CRMs at enhancer regions have similar function diversity. However, currently, very limited high-resolution chromosome interaction data, such as ChIA-PET, is available so that the mapping between enhancers and genes still has a lot of ambiguity, especially considering that some enhancers can regulate multiple target genes, none of which may be located nearby.

Here we mainly applied ChIP-GSM to cell type-specific data, as most ChIP-seq profiles are generated using specific cell lines. ChIP-GSM can also be applied to ChIP-seq data sequenced from human tissues, for which the binding signals can be much noisier and exhibit more variable binding strengths than in vitro cell lines. Applying ChIP-GSM to those cases has the potential to uncover more detailed TF-associations not well captured by conventional methods, thereby leading to new biological insights relevant to human disease.

## 4. Methods

### 4.1 ChIP-GSM: cis-regulatory module inference

The computational complexity of CRM inference (studying associations between TFs) increases exponentially with the number of TFs (*T*). Therefore, for a large set of TFs, exploring exhaustively all possible combinations becomes impractical (e.g., with 2^50^ TF combinations for *T* = 50). To control the searching scale of CRMs, ChIP-GSM first identifies a list of candidate CRMs based on ChIP-seq read count observations at regulatory regions (***Supplementary Fig. S1***) and constructs a matrix **B** with *M* rows (the total number of candidate CRMs) and *T* columns (the total number of TFs), where *M* ≪ 2^*T*^. In this matrix, each row is a binary vector [*b*(*m*,1),*b*(*m*,2),…,*b*(*m,t*),…,*b*(*m,T*)] representing a unique CRM. For modelling background regions that are not regulated by any CRMs, we add an additional all-zero row (*m* = 0) to **B**.

Given *K* regions, we further define a CRM index variable *c*_*k*_ for each specific region *k*. *c*_*k*_ can take value from 0 to *M*, indexing a candidate CRM (a row in matrix **B**). We control that there is at most one non-zero module regulating each individual region in each round of sampling so we have *c*_*k*_ ∈ [1, *M*] if this region is regulated by a CRM or *c*_*k*_ = 0 otherwise. As each unit in matrix **B** is binary, if *b*(*c*_*k*_, *t*) = 1, region *k* will be treated as a binding region of TF *t*; or *c*_*k,m*_*b*_*m,t*_ = 0 and region *k* is a background region. The observed read count of TF *t* at region *k, Y*_*k,t*_, is modeled using a mixture model as follows:

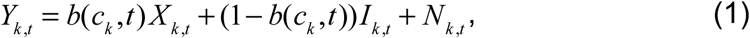

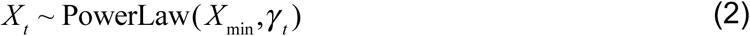

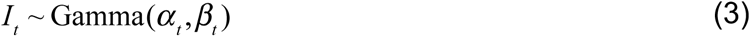

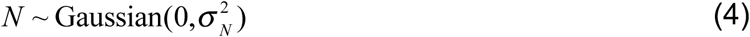

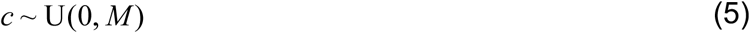

Where *X*_*k,t*_ represents the read count of a TF binding region and is assumed to follow a Power-Law distribution with hyper-parameters *X*_min_ and *γ*_*t*_; *I*_*k,t*_ represents the read count if the region is a background region and is assumed to follow a Gamma distribution with mean and shape parameters *α*_*t*_ and *β*_*t*_; and *N*_*k,t*_ represents the read count noise in the ChIP-seq data and is assumed to follow a zero-mean Gaussian distribution with variance 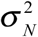. On the variable 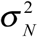, we assume a prior Inverse-Gamma distribution (conjugate prior of the variance of Gaussian distribution) with mean and shape parameters *α*_*N*_ and *β*_*N*_.

Specific for gene promoters, as demonstrated in [29, 30], there is an exponential decay effect on read enrichment along with the increase of relative distance to nearest transcription starting site (TSS). At background regions, the distribution of read enrichment is usually uniform. For enhancer studies, because both binding and background regions are distal to TSSs, the regulatory effects are independent of their binding locations considering the loop structure between enhancers and target genes.

To better facilitate identifying regulatory regions with TF bindings at promoter area, we model the effects of a regulatory region on target gene using a distance-based mixture model as follows:

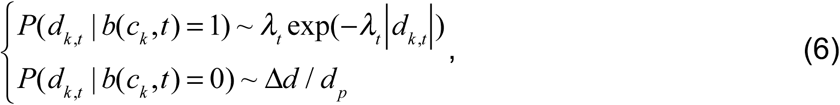

where *d*_*k*_ represents the relative distance of region *k* to the nearest TSS; *λ*_*t*_ is the exponential decaying parameter; Δ*d* represents the region length (500 bps); and *d*_*p*_ represents the length of promoter area around TSS (20k bps: +/− 10k bps around TSS). *λ*_*t*_ is TF-specific and unknown. It needs to be estimated from the relative distance distribution of binding regions of TF *t*.

Given observed ChIP-seq read counts **Y**, candidate modules **B**, and the relative binding locations to TSS **D**, we jointly and iteratively estimate CRM indexes **C** for all regions, read counts **X** at bindings regions or **I** at background regions for all TFs in a probabilistic framework as follows:

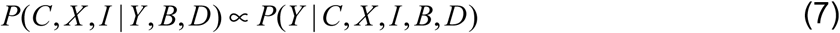

Variables of **C, X** and **I** are dependent as for each region, the binding status of each TF is deteremined by the CRM. Simutaniously estimating a joint probability distribution for all variables is very chanllenging. The main purpose is to estimate the distribution of CRMs. Therefore, we use Gibbs Sampling technic to sample variables iteratively as follows and finally use CRM samples to appromate its posterior probability distirbution.

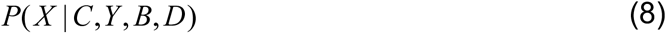

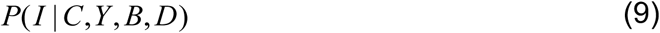

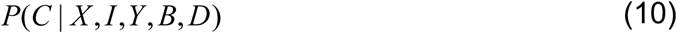

#### Model initialization

To initiate the model, ChIP-GSM first calculates *T* weights for each region, corresponding to the binding likelihoods of *T* TFs, where each weight is estimated based on the TF read count *X*_*k,t*_ (or *I*_*k,t*_) given the binding or background status of region *k*. For TF *t*, we observed totally *R*_*t*_ reads across all *K* regions from its ChIP-seq profile. To calculate the initial weight of each region, ChIP-GSM first roughly estimates the number of reads to be respectively assigned to binding and background regions by simulating the ChIP-seq sequencing process [6]. We assume that all regions are background regions and calculate weight *p*_*k,t*_ according to the observed read count.

We then select regions with read count *Y*_*k,t*_ >50 as TF binding regions and amplify their weight by *F* times. The number of reads aligned to binding regions can then be estimated as:

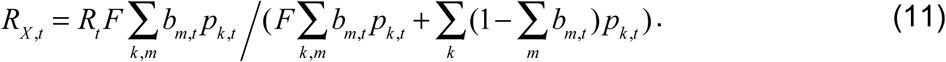

We assign *R*_*X, t*_ reads one by one to each of its targeting regions according to their amplified weights and get an initial read count *X*_*k,t*_ for each. Similarly, we assign remaining reads (*R*_*t*_ -*R*_*X, t*_) to each of the other background regions accoding to the initial weights and get read count *I*_*k,t*_. In general, *X*_*k,t*_ or *I*_*k,t*_ will differ from the observed ChIP-seq read count *Y*_*k,t*_ for the same location. We aim to minimize this difference globally at all regions and jointly for all TFs by iteratively estimating CRMs, which control the binding states of all TFs at each location.

#### Sampling ChIP-seq read counts

To estimate the read count for each region, ChIP-GSM updates the weight of every region, assigns reads to them probabilistically and then obtains a new read count for each region. Specifically, given the number of reads assigned to each region, region binding state, and the observed ChIP-seq read count, ChIP-GSM calculates a conditional probability as that region’s updated weight:

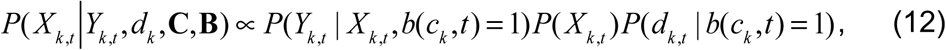

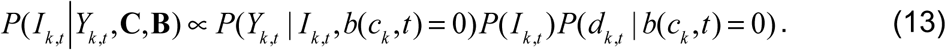

Then for a binding region, according to in the PowerLaw distribution, it must contains at least *X*_min_ reads. Thus, ChIP-GSM assigns *X*_min_ reads evenly to all binding regions and then assigns the remaining reads 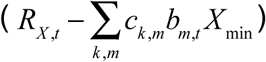 one by one to each of them according to the distribution of their updated weights. Finally, each binding region has a new read count *X*′_*k,t*_. Similarly, for background regions, ChIP-GSM assigns *R*_*t*_ -*R*_*X, t*_ reads one by one to them according to the distribution of updated weights and each background region has new a read count *I*′_*k,t*_. Finally, for every region, an estimated read count *Y*′_*k,t*_ (*X*′_*k,t*_ *b*(*c*_*k*_, *t*) + *I*′_*k,t*_ (1− *b*(*c*_*k*_, *t*))) is generated, with a read count difference *N*_*k,t*_ from the observed read count *Y*_*k,t*_. We control the variance 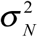 between **Y**′ and **Y** across all regions for *T* TFs using a Gaussian disrtibution. The conditional probability of 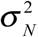 is calculated as follows:

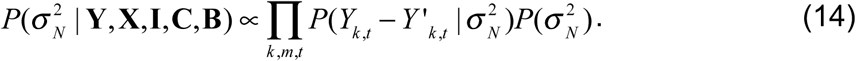

As detailed in ***Supplementary Section S2***, Eq. (9) is an inverse-Gamma distribution so we can directly sample 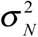 with updated mean *α*_*N*_ + *KT* / 2 and shape parameter 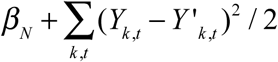.

#### IV. Sampling CRMs

To sample a CRM for each region from all candidate CRMs, ChIP-GSM estimates a discrete probability density distribution. Specifically, for each region *k* and candidate CRM *m*, we calculate a conditional probability, and sample a CRM for *k* accordingly (see ***Supplementary Section 2*** for details).

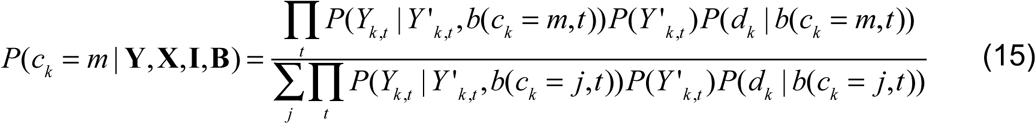

After repeating CRM sampling at all regions, we enter an updated matrix **C** into Eq. (6) to get a updated list of foreground and background regions and also the corresponding total numbers of reads for each category, from which we start a new round of sampling based on Eq. (7) and (8). We run the sampling process until the sampler appears to converge on the equilibrium distribution and then start accumulating samples on CRMs. After drawing enough samples, we obtain a weighted matrix **Ĉ** with each element 0 ≤ *ĉ*_*k,m*_ ≤ 1 denoting the sampling frequency of CRM *m* for region *k*. *ĉ*_*k,m*_ provides a posterior probability of module *m* regulation of region *k*. Although in each round of sampling, we sample only one *c*_*k,m*_ = 1 for region *k*, after drawing many samples, in **Ĉ** there will exist a probabilistic distribution of all candidate CRMs for *k*, with multiple *ĉ*_*k,m*_ > 0 being highly likely. Such information indicates how TFs associate with each other and estimates the strength of each association. More detailed sampling procedure is provided in **Supplementary Methods**.

### 4.2 ChIP-GSM: CRM-based active regulatory region prediction

We aim to use the weighted matrix **Ĉ** to further investigate active regulatory regions under the assumption that a foreground region bound by multiple TFs is likely (though not necessarily) an active regulatory element. By combining inferred CRMs probabilities with experimentally measured activities for regulatory regions in the context of a specific cell type, we can train a classifier (i.e., determine an optimal set of weights ***β*** for CRMs) to predict the activity of every element bound by CRMs. To weight the identified CRMs in this way, we train a binomial model using elastic net logistic regression [31], which is a natural fit for this application because the CRMs are highly correlated (sharing TFs) and have strong grouping effects. Elastic net regression assigns similar weights to correlated features or removes them altogether by assigning zero weights [7]. Unlike linear regression, elastic net regression extends the method of least squares by adding a regularization (or penalty) that includes the weights ***β*** in the minimization process:

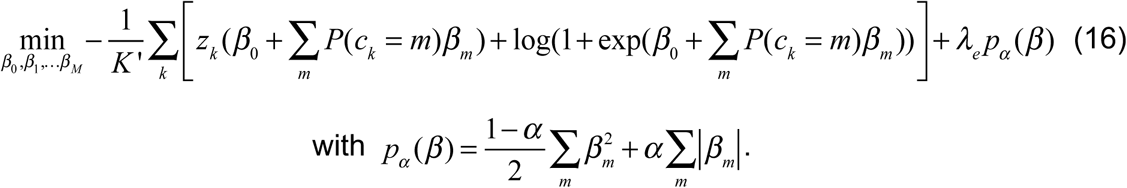

where *z*_*k*_ is a binary (+/−) label for regions with experimental cell-type specific activity measurments; *K*′ is the total number of labelled positive or negative regions; *λ*_*e*_ ≥ 0 is a parameter controlling the model complexity; 0 ≤ *α* ≤ 1 controls the relative contributions of ridge regression and LASSO to the overall regularization penalty. After the training, we obtain an optimal set of weights including *β*_0_ and *β*_1_… *β*_*m*_ … *β*_*M*_ for individual CRMs.

### 4.3 Performance evaluation with realistic simulation data

We simulated ChIP-seq read counts derived from real ChIP-seq data of 20 TFs, downloaded from ENCODE database (http://genome.ucsc.edu/ENCODE/). The advantages of this design are: (1) in real data, each ChIP-seq profile is unique in both ChIP-seq read count distribution and binding locations of a specific TF; (2) strong and weak binding events are both included, as TFs have different binding strengths; and (3) real associations between TFs are retained. We performed three simulations with four TFs in Case #1, seven TFs in Case #2, and 20 TFs in Case #3 (read count distributions are provided in ***Supplementary Fig. S8***). In each case, we generated 10 replicates by randomly shuffling read counts at different locations. We compared ChIP-GSM with competing methods jMOSAiCS, and SignalSpider. Each method’s performance is based on the recovery of all ‘true’ bindings simultaneously at all regions using the F-measure (F=2/(1/precision+1/recall)) either of all modules (***Supplementary Fig. S9A***) or of modules containing at least one ‘weak’ binding (read count < 30) (***Supplementary Fig. S9B***). In all cases, ChIP-GSM provides improved performance over the existing comparable methods.

### 4.4 Benchmarking ChIP-GSM on modules with known associations

For benchmarking we used a H1-hESC cell ChIP-seq dataset with factors EZH2, SUZ12, H3K27me3 and H3K4me3 from the ENCODE database. Associations among these four factors have been well demonstrated in previous biological studies: EZH2 and SUZ12 are Polycomb-group (PcG) proteins [32] and tend to co-bind to ‘bivalent’ domains marked by H3K27me3 and H3K4me3 [33, 34]. Given these strong associations, a large number of regions co-activated by all four factors are expected. Applying ChIP-GSM to this dataset we identified 6,564 regions (29%) regulated by all four factors, while SignalSpider and jMOSAICs identified much less, 21.6% and 9.65% of such regions, respectively.

### 4.5 Enrichment of histone modifications and ChIA-PET interactions

Cell type-specific ChIP-seq peaks of H3K27ac, H3K4me1 and H3K4me3. All genomic coordinates were referred to hg19. H3K27ac and H3K4me1 are widely used as markers for distal enhancers while H3K4me3 is more often used as a marker for proximal promoters [21]. For each histone marker, we label each region ‘+’ if the region center is within 2k bps from any peak centers; or we label it as ‘-’. For each cell type, two categories of regions were extracted as (H3K27ac ‘+’, H3K4me1 ‘+’, H3K4me3 ‘-’) and (H3K27ac ‘-’, H3K4me1 ‘-’, H3K4me3 ‘+’). Hypergeometric p-value was calculated for the enrichment of labeled regions for each category within top 10% ChIP-GSM predicted cell type-specific enhancers or promoters, compared to the enrichment situation among all enhancers or promoters. Cell type-specific ChIA-PET chromatin interactions were downloaded from the ENCODE database, too. To eliminate nonsense interactions, all interactions were first filtered using all enhancers and promoters and each retained interaction is a loop between an enhancer and a promoter. Then, for each enhancer or promoter, we count the number of loops around it and for all enhancers or promoters, Pearson correlation coefficient is calculated between ChIP-GSM predicted regulatory activity and the number of ChIA-PET loops.

### 4.6 K562 time-course gene expression

We downloaded a time-course K562 gene expression dataset [24] from the GEO database (GEO accession number: GSE1036). K562 cells (duplicate cultures A & B) are treated with 50 micromolar hemin for 0, 6, 12, 24, 48, 72 hours followed by RNA extraction and gene expression profiling on Affymetrix human U133A arrays. Under each time point there are two duplicates as A0 and B0 under time point ‘0’ and Ai and Bi under each time point ‘i’. For a pair of TFs, we calculate the Pearson correlation coefficient using their mRNA transcription, assuming associated TFs in the same module are more likely to be active at the same time, although relationship between the protein activity and mRNA transcription may not be linear. We also select up or down-regulated genes if at any time point *i*, compared to time point ‘0’, the gene expression log2 fold change is larger than 1: |log2(A0)-log2(Ai)|>1, |log2(A0)-log2(Bi)|>1, |log2(B0)- log2(Ai)|>1, and |log2(B0)-log2(Bi)|>1. In total, we collect 1,686 genes and further cluster them into eight clusters using hierarchical clustering (**Fig 4D**).

## Code access

R scripts of ChIP-GSM and its user manual are made available to the research community at http://www.cbil.ece.vt.edu/software.htm.

## Acknowledgments

This work was supported in part by the National Institutes of Health (CA149653 to J.X., CA149147 & CA184902 to R.C., CA164384 to L,H.-C., and GM125878 to A.F.N).

