## Supplementary figures and methods for "Inferring active cis-regulatory modules to predict functional regulatory elements"

### Supplementary Results

#### S1. Workflow of ChIP-GSM

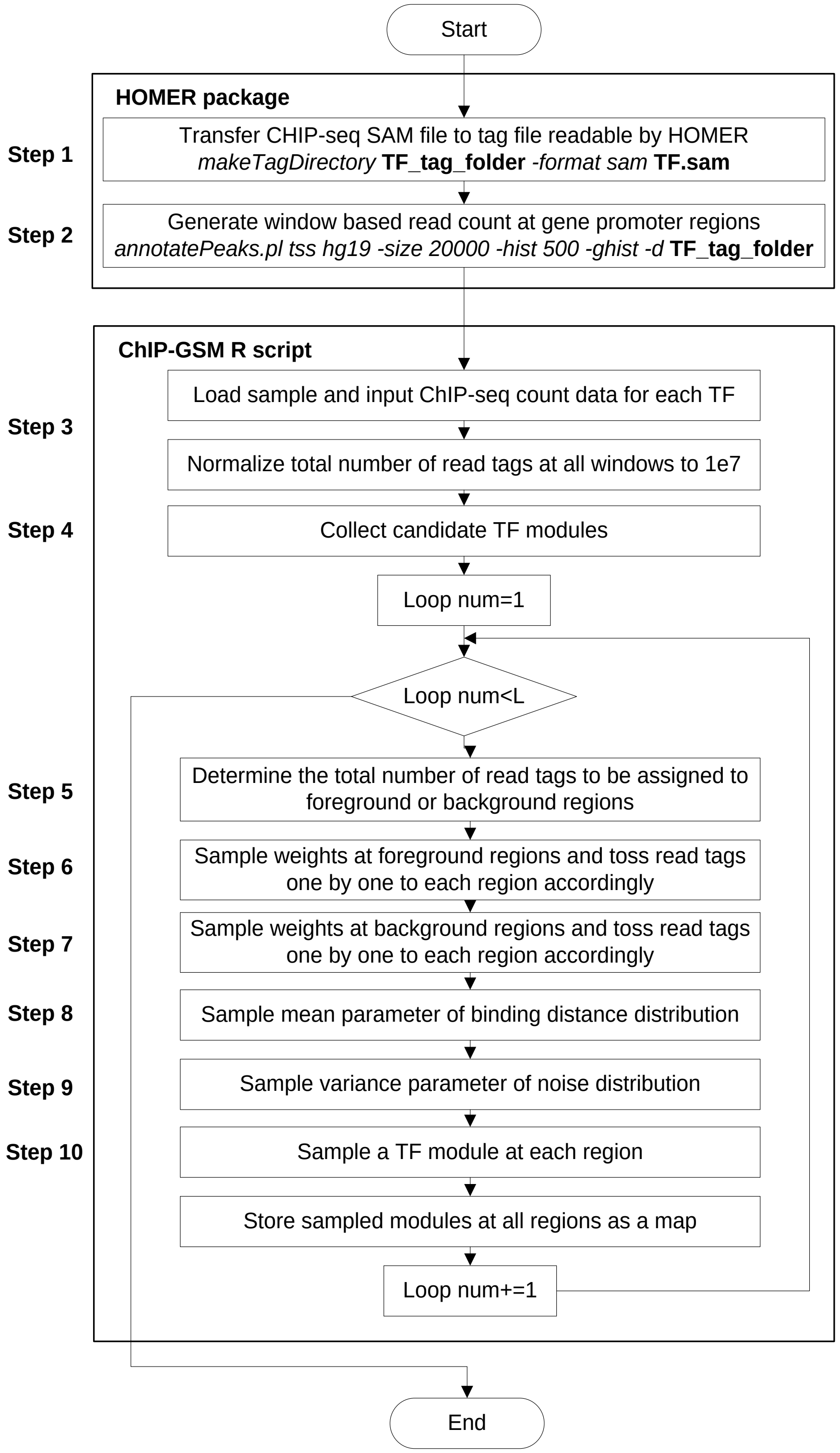

**Fig. S1.** Workflow of the ChIP-GSM approach.

#### ChIP-seq data pre-processing

**Step 1**: We use HOMER (v4.9) to process ChIP-seq read count profiles (BAM files; aligned to human reference genome hg19) into read tag files where 5' start locations as well as directions of read tags in each chromosome are stored in an individual file.

**Step 2**: We partition human cell type-specific enhancer or promoter-like regions (referred to hg19; downloaded from ENCODE webserver http://screen.encodeproject.org/) into 500 bps segments (regions less than 500 bps will be round up to 500 bps around the original region center) and count reads uniquely aligned to each segment.

**Step 3**: We normalize the total number of reads in each ChIP-seq profile to $10^7$. This step is necessary and important to eliminate any bias caused by the sequencing depth when we pool multiple ChIP-seq profiles for a joint analysis.

**Step 4**: ChIP-GSM can either take a list of pre-generated cis-regulatory modules (CRMs) as input or automatically identify a set of candidate CRMs based on the input ChIP-seq data.

#### Cis-regulatory module inference using ChIP-GSM

**Step 5**: We initiate the model by assigning possible CRM structures to each region. Then, based on the read counts, we determine regions potentially with or without bindings of each TF. For example, a region with more than 10 reads or 2 folds to the control ChIP-seq profile is likely to contain a binding site (a foreground region); otherwise it is a background region. We roughly estimate the total number of reads sequenced from foreground or background regions. After that, we process foreground and background regions separately as we assume their read counts follow different distributions.

**Step 6**: For each TF, we calculate a weight for each foreground region based on its read count, by assuming a Power-Law distribution. Here, the Power-Law distribution parameters are TF-specific, which can be obtained from distribution fitting of read counts in each TF ChIP-seq profile. We assign reads (the total number is determined in **Step 5)** one by one to all foreground regions according to their weights.

**Step 7**: For the same TF but background regions, we calculate a weight for each based on its read count and a Gamma distribution assumption. Parameters of this Gamma distribution are obtained by fitting read counts of the control ChIP-seq profile. We assign the remaining read tags in current TF ChIP-seq profile one by one to background regions according to their weights.

**Step 8**: Specially for promoter study, for each TF, we estimate the mean parameter for the Exponential distribution modelling the relative distance of foreground regions to the nearest transcription starting site (TSS), using their genomic locations. **Steps 5~8** are repeated for each TF.

**Step 9:** We estimate the variance of the residuals between observed and assigned read counts across all segments of all TFs.

**Step 10:** For each region, we calculate a conditional probability for each candidate CRM given the assigned read counts and randomly select a CRM according to their conditional probability distribution. This current region is classified as a foreground region for TFs within the sampled CRM or a background region. Results of **Step 10** are recorded and then brought back to **Step 5** to start the next round of sampling.

We run the sampling process until the sampler appears to converge on the equilibrium distribution and then start accumulating samples on CRMs. After collecting enough samples, the sampling frequency of each module-region unit denotes the posterior probability for binding occurrence. Foreground regions regulated by each module can be identified.

#### S2. CRM inference comparison using ChIP-seq data of K562 cells

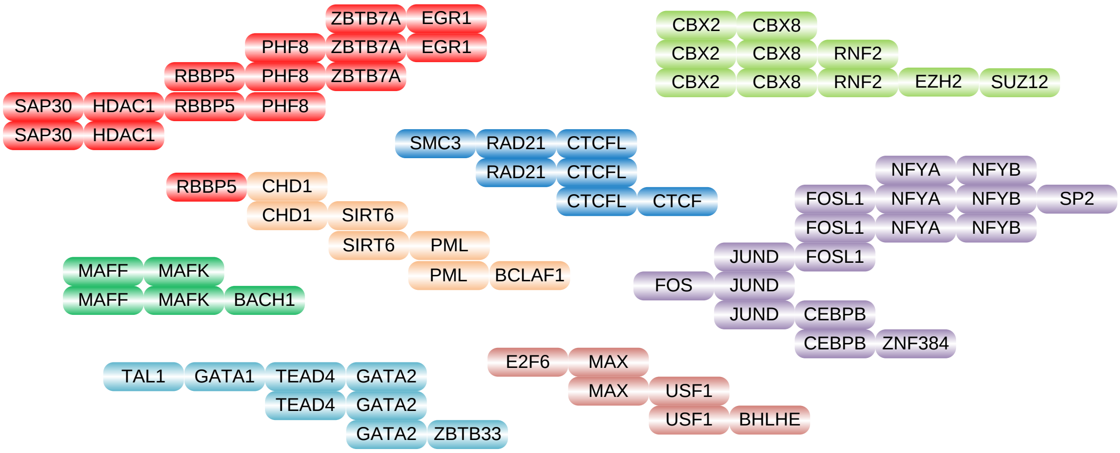

**Fig. S2.** Top 30 ChIP-GSM inferred CRMs from K562 promoter regions.

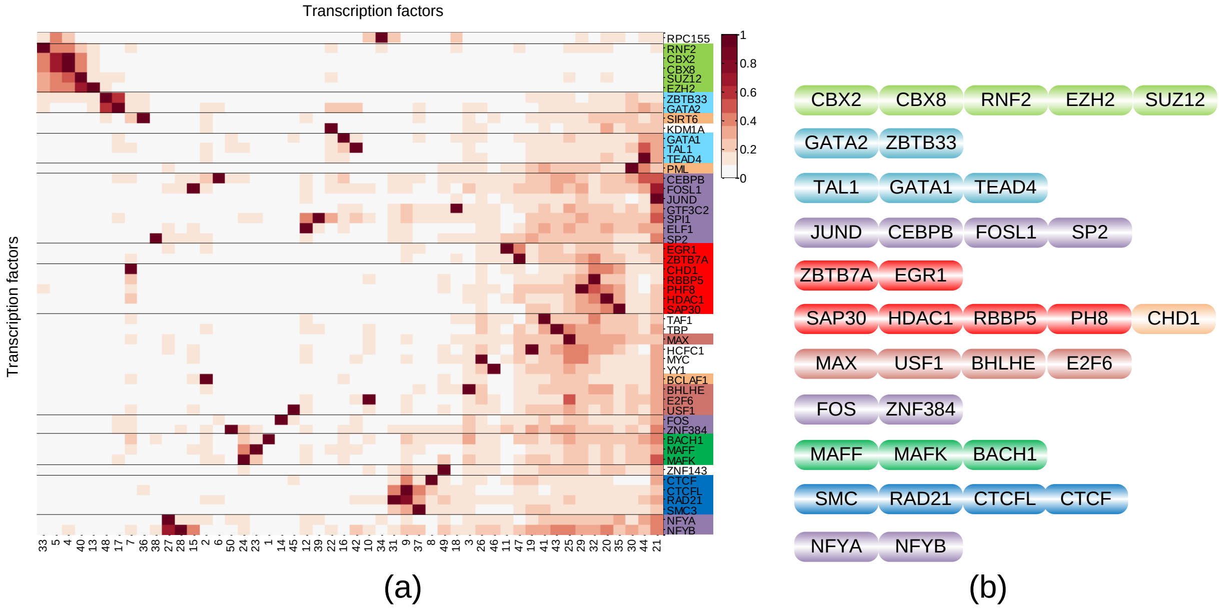

**Fig. S3. Inferred CRMs using the Rulefit approach** (1). (a) Hierarchical clustering of relative importance (RI) matrix estimated by Rulefit. (b) 11 inferred CRMs using the same group color as ChIP-GSM, but not able to predict more specific associations of TFs.

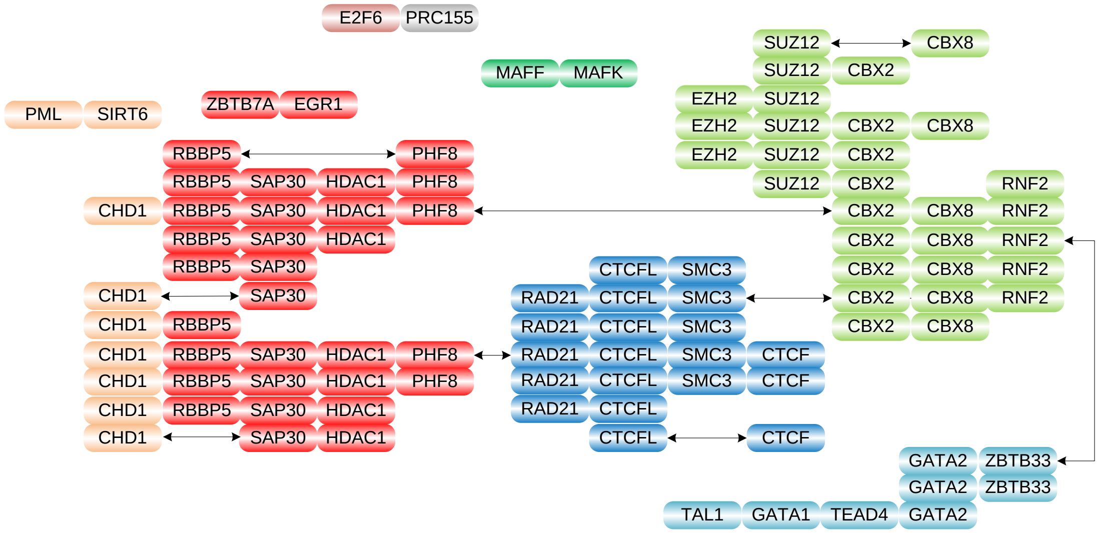

**Fig. S4. CRMs inferred by ISA** (2). CRMs predicted by ISA can be roughly clustered into four major groups (similar to the Groups 1, 2, 4 and 6 inferred by ChIP-GSM). However, CRMs in Groups 5 and 8 identified by ChIP-GSM are missing in the results of ISA.

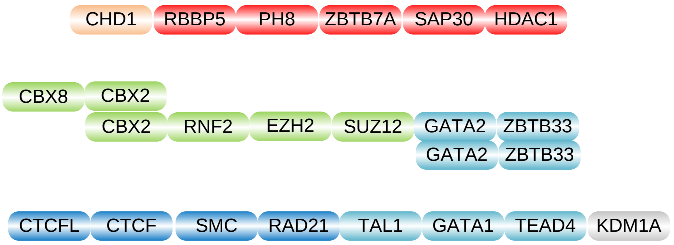

**Fig. S5. CRMs inferred by Plaid** (3). Plaid identified five modules with only high-level large-scale associations captured.

#### S3. K562 TF expression and functions

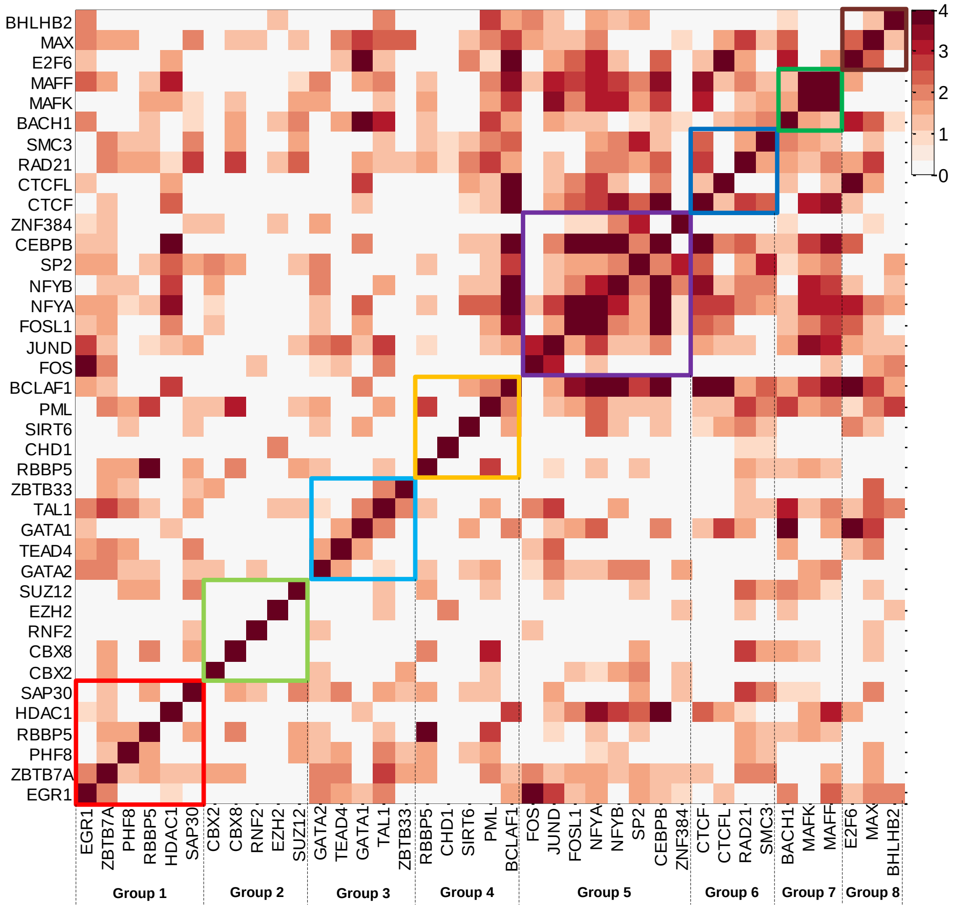

**Fig. S6. Co-expression of TFs in inferred CRMs**. Gene expression data was downloaded from GEO database with ID: GSE1036. Color bar represents –log10(*p*-value) of Pearson correlation coefficient. Rectangles with different colors represent the ChIP-GSM identified CRM groups.

#### TF functions:

In Group 1, the similarity for TFs is very significant (red blocks). Two associations, RBBP5-PHF8-SAP30-HDAC1 and EGR1-ZBTB7A, have been validated in K562 cells (1,4). A novel association PHF8–ZBTB7A is predicted in CRMs 9 and 37; the *p*-value of the mRNA expression correlation is 3.75e-2.

In Group 2, all TFs (i.e., CBX2, CBX8, RNF2, EZH2 and SUZ12) are Polycomb-group proteins whose associations have been established under stem cells and further verified to be existent in K562 cells (5). However, their expression patterns are quite diverse so that there is no significant correlation on their expression patterns.

In Group 3, pairwise correlation is highlighted but there is no significant triple wise correlation. One association GATA1-GATA2-TAL1 has been previously observed in (1) based on whole genome study. Due to existing evidence in support of the associations of GATA3-GATA2 and GATA3-TEAD4 (6,7), a high-order association of GATA2-GATA3-TEAD4 can be expected. However, due to the lack of GATA3 ChIP-seq data, only GATA2-TEAD4 is identified; the *p*-value of expression correlation is 3.29e-2. GATA2-ZBTB33 is confirmed by observing binding signals enrichment of GATA2 at regions regulated by ZBTB33 in K562 cells (8).

In Group 4, the combinations of TFs are more diverse and the supporting evidence of mRNA expression correlation is weak. The co-binding of RBBP5 and CHD1 has been observed at active promoters with enrichment of H3K4me3 (9,10). SIRT6 is also highly enriched at active promoters as well and its association with CHD1 is related to productive initiation (5).

Group 5 is a large group where two high-order associations, SP2-NFYA-NFYB and FOSL1-FOS-JUND-CEBPB, have been observed, which are also validated by another study on K562 cells (11). The overall correlation of mRNA expression of Group 5 TFs is very strong, as dense dark red units are observed in the Group 5 block. Here, a novel association, FOSL1-NFYA-NFYB, is also predicted, which is strongly supported by their co-binding enrichment at promoter regions of a large number of genes as well as their significant mRNA expression correlation (the core dark red block in Group 5).

In Group 6, the association of RAD21-SMC3-CTCF can be verified based on fact that both RAD21 and SMC3 are members of the cohesin complex, which is known to interact with CTCF (12) and act as an insulator (13). Their triple-wise mRNA co-expression correlation can be predicted based on their highly significant pairwise correlation relationship. Since CTCFL is a paralog of CTCF, its interaction with RAD21 or SMC3 is also supported by their binding signal enrichment at the same regions.

Group 7 only has three TFs but the association of BACH1 with the MAF protein complex including MAFF and MAFK is very strong, especially with MAFK to form a heterodimer (14). As shown in Fig. S5, BACH1 has a strong correlation with either MAFF or MAFK.

In Group 8, E2F6 may be recruited by MAX to gene promoters via E boxes (CACGTG) via protein-protein interaction (15). MAX, USF1, and BHLHE40 (BHLHB2) are E-box binding factors so that two associations of them including MAX-USF1 and BHLHE40-USF1 are identified by the ChIP-GSM approach. The correlation between E2F6 and MAX is very high. Since MAX and BHLHE40 are indirectly connected with USF1 (the expression profile of which is not available in our study), the correlation of MAX and BHLHE40 is weaker but still significant.

#### S4. Genes regulated by ChIP-GSM inferred CRMs for K562 promoter

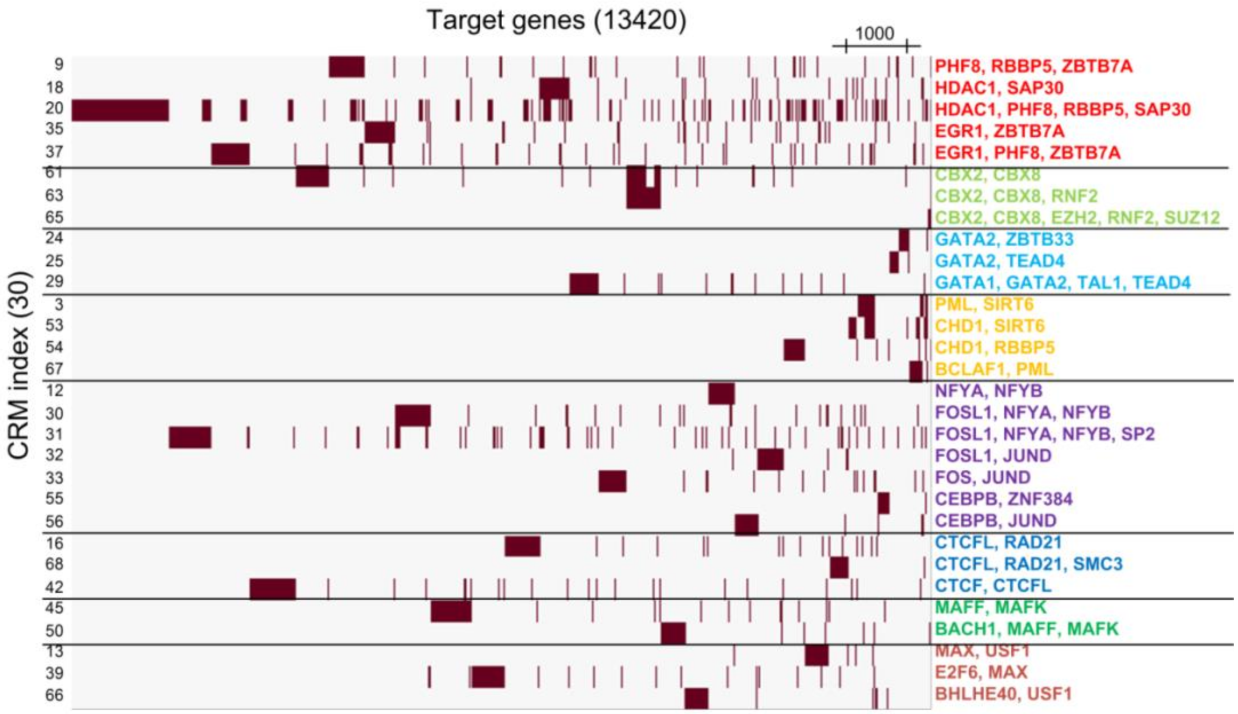

**Fig. S7. Target genes regulated by selected 30 CRMs functioning at K562 promoter.**

#### S5. ChIP-seq data simulation

Breast cancer MCF-7 cell line

**Case 1**: ER-α, GATA3, SIN3A, NR2F2

**Case 2**: CTCF, E2F1, MBD3, NR2F2, PML, POLR2A, RAD21

**Case 3**: CEBPB, CREB1, CTCF, E2F1, EP300, ER-α, FOXA1, GATA3, MBD3, MYC, NR2F2, PBX1, PML, POLR2A, RAD21, SIN3A, TCF7, TCF12, TLE3, ZNF217

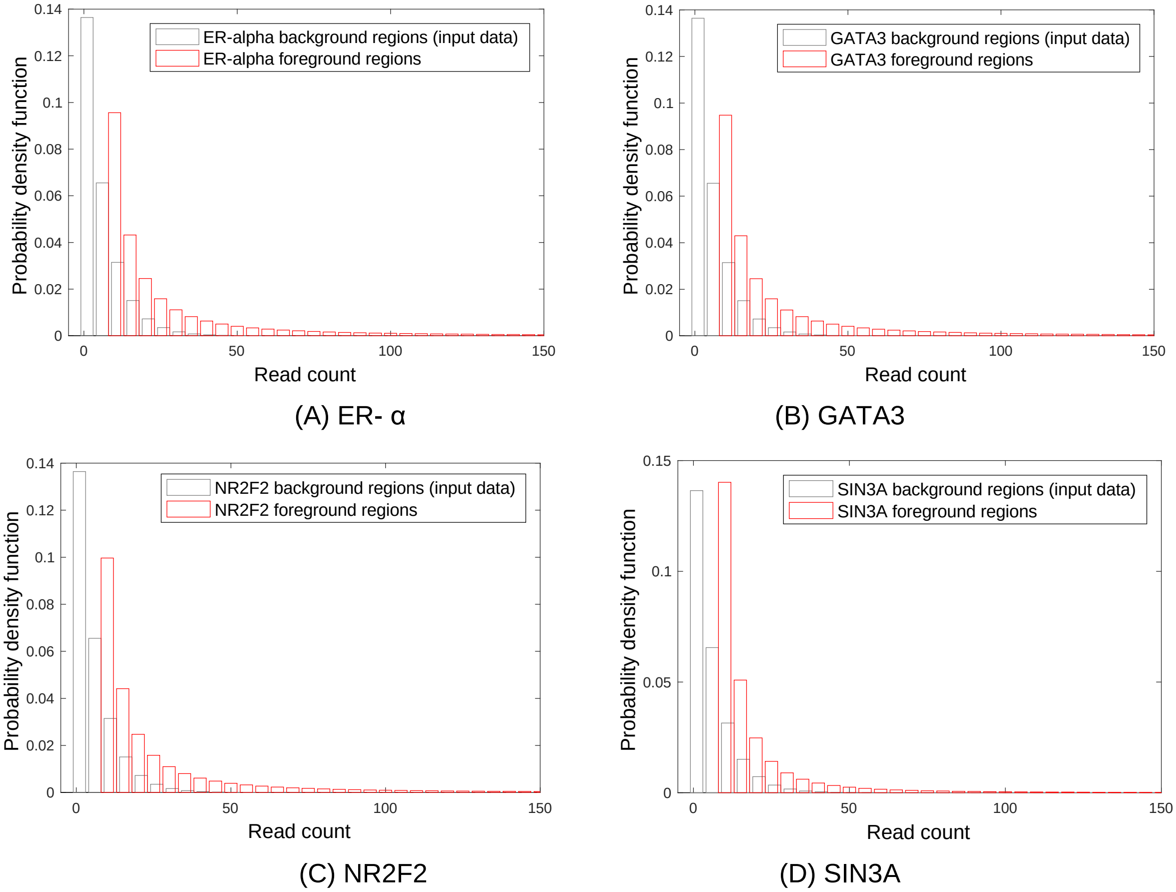

**Fig. S8. Distributions of simulated ChIP-seq read counts for Case 1**. For each TF, we count the TF ChIP-seq read tags falling in each region (500 bps around peak summit). We also count read tags in the matched control ChIP-seq profile. Repeating the pre-processing for every TF, we select regions with non-zero read counts of at least two TFs. Then, among these regions, for each TF, we randomly and respectively perturb the read counts at binding or non-binding regions. For each case, we generate 10 replicates.

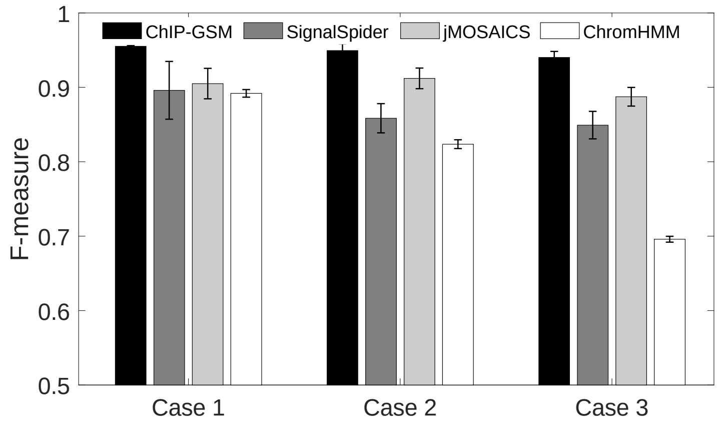

(A)

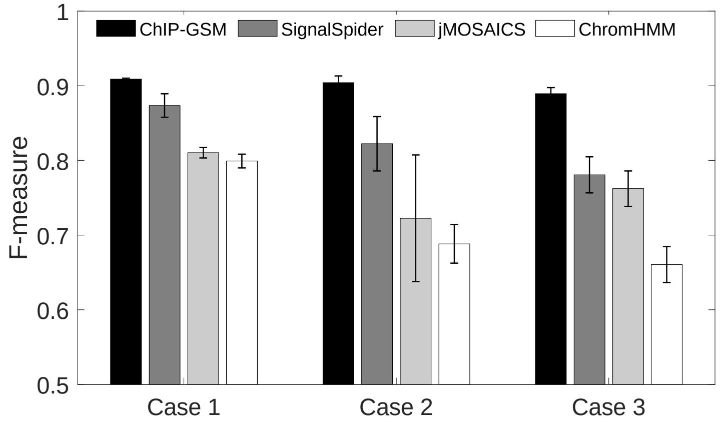

(B)

**Figure S9. Comparing ChIP-GSM against competing methods on CRM inference using realistically simulated ChIP-seq data.** (A) F-measures of each method on all regions; (B) F-measure of each method on regions with weak binding events. SignalSpider and jMOSAICS were developed to infer CRMs using binding signals of a small number of TFs. ChromHMM was developed to identify chromatin state using histone modifications. ChIP-GSM provides improved performance especially when there are a large number TFs with weak binding events.

#### Supplementary Methods

$X_{k,t}$ (read count at foreground region with $b(c_k,t)=1$) follows a Power-Law distribution:

$$P(X_{k,t}) \sim \frac{\gamma_t - 1}{X_{\min}} \left( \frac{X}{X_{\min}} \right)^{-\gamma_t}. \quad \text{(S-1)}$$

$I_{k,t}$ (read count at background region with $b(c_k,t)=0$) follows a Gamma distribution:

$$P(I_{k,t}) \sim (I)^{\alpha_{I,t}-1} \exp(-\beta_{I,t} I). \quad \text{(S-2)}$$

$N_{k,t}$ (read count noise at each region) follows a Gaussian distribution:

$$P(N_{k,t}) = \frac{1}{\sqrt{2\pi\sigma_N^2}} \exp\left(-\frac{N^2}{2\sigma_N^2}\right). \quad \text{(S-3)}$$

$\sigma_N^2$ (noise variance at all regions) follows an Inverse Gamma distribution:

$$P(\sigma_N^2) = \frac{1}{C_N} (\sigma_N^2)^{-\alpha_N - 1} \exp\left(-\frac{\beta_N}{\sigma_N^2}\right). \quad \text{(S-4)}$$

$d_{k,t}$ (relative distance of each region to the nearest TSS) follows Exponential distribution for $b(c_k,t)=1$ or Uniform distribution for $b(c_k,t)=0$:

$$\begin{cases} P(d_{k,t} \mid \sum_m c_{k,m} b_{m,t} = 1) \sim \lambda_t \exp(-\lambda_t |d|) \\ P(d_{k,t} \mid \sum_m c_{k,m} b_{m,t} = 0) \sim \text{Uniform}(d) \end{cases}, \quad \text{(S-5)}$$

where $\lambda_t$ (exponential factor) follows Gamma distribution as follows:

$$P(\lambda_t) = \frac{1}{C_\lambda} \lambda^{\alpha_\lambda - 1} \exp(-\lambda \beta_\lambda). \quad \text{(S-6)}$$

**Gibbs sampling procedure:**

**(1) Determine the total number of tags**

For the *t*-th TF, we randomly assign a weight $p_{k,t}$ to the *k*-th region according to Gamma distribution. Based on initial selection or previous round of sampling, we have a module-region regulatory map and know which module or TF is binding at which region, as $\mathbf{C}$. For any region with state $b(c_k,t)=1$, we amplify its weight by $F$ times. Then we can roughly estimate the total number of read tags assigned to foreground regions as follows:

$$R_{t,X} = R_t \frac{F\sum_k b(c_k,t)p_{k,t}}{F\sum_k b(c_k,t)p_{k,t} + \sum_k (1-b(c_k,t))p_{k,t}}, \quad \text{(S-7)}$$

where $R_t$ is the total number of read tags of the *t*-th TF, which is normalized to $10^7$ in this study. The total number of read tags $R_{t,I}$ assigned to background regions for the *t*-th TF can be calculated as $R_t - R_{t,X}$.

**(2) Sampling read counts for foreground regions**

Conditional probability of $X_{k,t}$ can be calculated as follows:

$$\begin{aligned} P(X \mid Y_{k,t}, d_k, b(c_k,t)=1, \Phi) &\propto P(Y_{k,t} \mid X, b(c_k,t)=1, \sigma_N^2)P(X)P(d_k \mid b(c_k,t)=1) \\ &\propto \frac{1}{\sigma_N}\exp\left[-\frac{(Y_{k,t}-X)^2}{2\sigma_N^2}\right]\cdot\frac{\gamma_t-1}{X_{\min}}\left(\frac{X}{X_{\min}}\right)^{-\gamma_t}\cdot\lambda_t\exp(-\lambda_t|d_k|) \end{aligned} \quad \text{(S-8)}$$

We calculate a weight $p_{k,t}$ for each region according to Eq. (S-8) as follows:

$$p_{k,t} = \frac{1}{C_{X,t}}\exp\left[-\frac{(Y_{k,t}-X'_{k,t})^2}{2\sigma_N^2}\right]\cdot\frac{\gamma_t-1}{X_{\min}}\left(\frac{X'_{k,t}}{X_{\min}}\right)^{-\gamma_t}\cdot\lambda_t\exp(-\lambda_t|d_k|), \quad \text{(S-9)}$$

where $C_{X,t} = \sum_k \exp\left[-\frac{(Y_{k,t}-X'_{k,t})^2}{2\sigma_N^2}\right]\cdot\frac{\gamma_t-1}{X_{\min}}\left(\frac{X'_{k,t}}{X_{\min}}\right)^{-\gamma_t}\cdot\lambda_t\exp(-\lambda|d_k|)$.

We first assign $X_{\min}$ reads to each foreground region to meet the requirement of PowerLaw distribution. Then we assign the remaining $R_{t,X} - \sum_k b(c_k,t) X_{\min}$ reads to all foreground regions according to the weight $p_{k,t}$ at each region and obtain a read count $X_{k,t}$ for each foreground region.

**(3) Sampling read counts for background regions**

Conditional probability of $I_{k,t}$ can be calculated as follows:

$$
\begin{aligned}
P(I \mid Y_{k,t}, b(c_k,t)=0, \Phi) &\propto P(Y_{k,t} \mid I, b(c_k,t)=0, \sigma_N^2) P(I) P(d_k \mid b(c_k,t)=0) \\
&\propto \frac{1}{\sigma_N} \exp\left[ - \frac{(Y_{k,t} - I)^2}{2\sigma_N^2} \right] \cdot \left[ (I)^{\alpha_{I,t}-1} \exp(-\beta_{I,t} I) \right] \cdot \frac{\Delta d}{d_p} \cdot
\end{aligned}
\qquad \text{(S-10)}
$$

We calculate a weight for each background region according to Eq. (S-10) and assign $R_{t,I}$ reads to background regions according to their weights. In detail, we randomly sample a temp count $I'_{k,t}$ based on Eq. (S-10) by varying $I$ from 1 to $X_{upper}$. Then, a weight $p_{k,t}$ at a background region can be calculated as:

$$
p_{k,t} = \frac{1}{C_{I,t}} \exp\left[ - \frac{(Y_{k,t} - I'_{k,t})^2}{2\sigma_N^2} \right] \cdot \left[ (I'_{k,t})^{\alpha_{I,t}-1} \exp(-\beta_{I,t} I'_{k,t}) \right] \quad , \qquad \text{(S-11)}
$$

where $C_{I,t} = \sum_k \exp\left[ - \frac{(Y_{k,t} - I'_{k,t})^2}{2\sigma_N^2} \right] \cdot (I'_{k,t})^{\alpha_{I,t}-1} \exp(-\beta_{I,t} I'_{k,t})$.

We assign $R_{t,I}$ reads to all background regions according to the weight $p_{k,t}$ at each region and finally obtain a read count $I_{k,t}$.

**(4) Sampling exponential parameter of relative distance**

Conditional probability of $\lambda_t$ can be calculated as follows:

$$P(\lambda_t \mid \mathbf{D},\mathbf{C},\Phi) \propto \prod_k P(d_k \mid b(c_k,t)=1)P(\lambda_t)$$
$$\propto \prod_{k|b(c_k,t)=1} \exp(-\lambda_t |d_k|)\lambda_t^{\alpha_{\lambda,t}-1}\lambda_t \exp(-\lambda_t \beta_{\lambda,t}) \quad . \qquad \text{(S-12)}$$
$$\propto \lambda_t^{\alpha_{\lambda,t}-1+\sum_k b(c_k,t)} \exp(-\lambda_t(\beta_{\lambda,t} + \sum_k b(c_k,t)|d_k|))$$

It can be seen from the form of the equation that Eq. (S-12) is also a Gamma distribution.

**(5) Sampling variance of noise component**

After repeating the steps according to Eq. (S-7) ~ (S-12) for all TFs, the conditional probability of $\sigma_N^2$ can be calculated as follows:

$$P(\sigma_N^2 \mid \mathbf{Y},\mathbf{X},\mathbf{I},\mathbf{C}) \propto \prod_k \prod_t P(Y_{k,t} \mid Y'_{k,t},\sigma_N^2)P(\sigma_N^2)$$
$$\propto \prod_k \prod_t \frac{1}{\sigma_N} \exp\left[-\frac{(Y_{k,t} - Y'_{k,t})}{2\sigma_N^2}\right] \sigma_N^{-2\alpha_N-2} \exp\left(-\beta_N \frac{1}{\sigma_N^2}\right) \quad .\text{(S-13)}$$
$$\propto \left(\sigma_N^2\right)^{-(\alpha_N+KT/2)-1} \exp\left(-\left[\beta_N + \frac{1}{2}\sum_k \sum_t (Y_{k,t} - Y'_{k,t})^2\right]\frac{1}{\sigma_N^2}\right)$$

It can be found that Eq. (S-13) is still an Inverse-gamma distribution.

**(6) Sampling CRMs for foreground regions**

For the *k*-th region, the conditional probability of each CRM can be calculated as follows:

$$P(c_{k,m} = 1 \mid \mathbf{Y},\mathbf{X},\mathbf{I},\mathbf{D},\mathbf{B},\Phi)$$
$$= \frac{1}{C_k}\left[\prod_{t|b(c_k,t)=1} P(Y_{k,t} \mid X_{k,t})P(X_{k,t})P(d_k)\right] \times \left[\prod_{t|b(c_k,t)=0} P(Y_{k,t} \mid I_{k,t})P(I_{k,t})P(d_k)\right] . \qquad \text{(S-14)}$$

where $C_k = \sum_m P(c_k = m \mid \mathbf{Y},\mathbf{X},\mathbf{I},\mathbf{D},\mathbf{B},\Phi)$.

For each region, we sample a CRM according to its conditional probability distribution and update the binding state of each region.

The MATLAB version of Elastic Net Logistic Regression package can be downloaded from https://web.stanford.edu/~hastie/glmnet_matlab/.

Parameter settings are listed as follows:

```
options.alpha = 0.1;
options.nlambda = 100;
options.standardize = true;
options.intr = true;
options.thresh = 1e-7;
options.cl  = [-Inf;Inf];
options.maxit = 1e+5;
options.ltype = 'Newton';
options.standardize_resp = false;
options.mtype = 'ungrouped';
family = 'binomial';
```
